# Cytosolic bacterial pathogens activate TLR pathways in tumors that synergistically enhance STING agonist cancer therapies

**DOI:** 10.1101/2024.01.30.578087

**Authors:** Meggie Danielson, Chris J. Nicolai, Thaomy T. Vo, Natalie Wolf, Thomas P. Burke

## Abstract

Bacterial pathogens that invade the eukaryotic cytosol are distinctive tools for fighting cancer, as they preferentially target tumors and can deliver cancer antigens to MHC-I. Cytosolic bacterial pathogens have undergone extensive preclinical development and human clinical trials, yet the molecular mechanisms by which they are detected by innate immunity in tumors is unclear. We report that intratumoral delivery of phylogenetically distinct cytosolic pathogens, including *Listeria, Rickettsia,* and *Burkholderia* species, elicited anti-tumor responses in established, poorly immunogenic melanoma and lymphoma in mice. We were surprised to observe that although the bacteria required entry to the cytosol, the anti-tumor responses were largely independent of the cytosolic sensors cGAS/STING and instead required TLR signaling. Combining pathogens with TLR agonists did not enhance anti-tumor efficacy, while combinations with STING agonists elicited profound, synergistic anti-tumor effects with complete responses in >80% of mice after a single dose. Small molecule TLR agonists also synergistically enhanced the anti-tumor activity of STING agonists. The anti-tumor effects were diminished in *Rag2*-deficient mice and upon CD8 T cell depletion. Mice cured from combination therapy developed immunity to cancer rechallenge that was superior to STING agonist monotherapy. Together, these data provide a framework for enhancing the efficacy of microbial cancer therapies and small molecule innate immune agonists, via the co-activation of STING and TLRs.

## Introduction

Bacteria that invade the eukaryotic cytosol are promising tools for treating cancer, as bacteria preferentially reside in tumors and can be engineered to deliver cancer antigens to MHC-I, eliciting potent CD8^+^ T cell responses^1–8^. Bacterial vaccine platforms have undergone extensive preclinical testing and human clinical trials^9–11^, however the contributions made by innate immunity to the anti-cancer response elicited by microbes are unclear. Activating innate immune receptors with small molecules in the tumor microenvironment (TME) elicits potent anti-tumor effects and has resulted in FDA-approval of anti-cancer drugs^12–15^, and therefore activation of these pathways by microbial vaccine platforms may contribute to their anti-cancer effects. Understanding the molecular mechanisms by which bacterial pathogens elicit anti-tumor responses will enhance our ability to design novel microbial and small molecule-based therapies for cancer immunotherapy.

Pattern recognition receptors (PRRs) detect pathogen-associated molecular patterns (PAMPs) and elicit pro-inflammatory cytokine responses that protect against infection^16,17^. Toll-like receptors (TLRs) are membrane bound PRRs that detect extracellular or endosomal microbial ligands. TLRs recruit cytosolic adaptors including MyD88 and TRIF to activate transcription factors including NF-κB, resulting in the secretion of pro-inflammatory cytokines such as tumor necrosis factor α (TNF-α)^18,19^. In contrast to membrane-bound TLRs, the protein cyclic GMP-AMP synthase (cGAS) binds mislocalized DNA in the cytosol as a signature of infection^20^. cGAS then synthesizes the cyclic dinucleotide (CDN) 2’3’ cyclic GMP-AMP (cGAMP), which binds to and activates stimulator of interferon genes (STING)^21–25^. STING activates Tank-binding kinase 1 (TBK1) and interferon responsive factor 3 (IRF3), causing a robust inflammatory response hallmarked by production of type I interferon (IFN-I), TNF-α, and chemokines^23,24,26–28^.

*Listeria monocytogenes* (*Lm*)*, Rickettsia parkeri* (*Rp*), and *Burkholderia thailandensis* (*Bt*) are three distantly related pathogens that share a similar intracellular lifecycle of replicating directly in the cytosol of mammalian cells. However, despite residing in the same cytosolic compartment, *Lm, Bt,* and *Rp* have distinct relationships with PRRs. *Lm* is a Gram-positive foodborne pathogen that activates STING via the secretion of the CDN cyclic-di-AMP^29,30^, and *Lm* also activates TLR2 and Myd88 *in vivo*^31–34^. In contrast, *Rp* is a Gram-negative tick-borne pathogen whose bacteriolysis can activate cGAS, but this activation is masked by inflammasome-mediated cell death^35^. Mice lacking the lipopolysaccharide receptor TLR4 have increased susceptibility to rickettsial infection, suggesting that *Rp* also activates TLRs *in vivo*^36,37^. *Bt* is a Gram-negative soil-dwelling microbe that is avirulent in humans, as it is strongly restricted by inflammasomes^38^ and is detected by TLRs^39^. Its interactions with cGAS/STING are uncharacterized. As cytosolic pathogens, these microbes have the capacity to deliver antigens to MHC-I, and *L. monocytogenes* has undergone human clinical trials as a cancer vaccine platform^1,9,11,40,41^, yet the underlying mechanisms by which *Lm, Rp* and *Bt* activate innate immunity in tumors are unknown.

Bacterial pathogens hold the potential to robustly activate innate immunity for cancer immunotherapy, and *Mycobacterium bovis* Bacillus Calmette-Guerinwhich (BCG) is approved for bladder cancer^42^. Activating innate immunity with small molecule TLR agonists has also been successful in the clinic, for example imiquimod targets TLR7 and is FDA-approved for basal cell carcinoma^43,44^. Intratumoral delivery of small molecule STING agonists potently inhibits tumor growth in preclinical models^12–14,45,46^. STING agonists activate CD8^+^ T cells and elicit long-lasting memory against cancer rechallenge^12,27,28^. However, human clinical trials using intratumoral delivery of STING agonists were not efficacious^47,48^, demonstrating the need for new approaches that enhance STING agonist therapies for cancer immunotherapy.

Here, we sought to determine how cytosolic bacterial pathogens activate innate immunity in tumors. We report that *Lm, Rp,* and *Bt* inhibited the growth of multiple, poorly immunogenic tumors in mice with no observable toxicity. We were surprised to find that the pathogens required cytosolic access for inducing anti-tumor effects, yet the anti-tumor activity was independent of cGAS/STING and instead required TLR signaling. The bacteria were more efficacious than small molecule TLR agonists and required IFN-I signaling. When combined with STING agonists, cytosolic pathogens elicited striking, synergistic anti-tumor effects and immunity to cancer cell rechallenge. Small molecule TLR agonists recapitulated synergy when combined with STING agonists. The combination therapy elicited long-lasting immunity against cancer cell rechallenge that required CD8^+^ T cells. Together, this study reveals underlying mechanisms by which microbes elicit anti-tumor responses and suggests that co-activation of STING and TLR pathways with microbes or small molecules elicits synergistic anti-tumor responses.

## Results

### Intratumoral delivery of cytosolic bacterial pathogens elicits dose-dependent anti-tumor responses in multiple non-immunogenic murine tumor models

It was unknown whether intratumoral delivery of cytosolic bacteria elicited anti-tumor responses and if intratumoral delivery caused toxicity *in vivo*. To limit any potential toxicity, we used attenuated λι*actA*λι*inlB* mutant *Lm* strains that underwent phase 1 and 2 clinical trials and are tolerated in humans at doses of >10^9^ bacteria^9,11,40,41^. This strain is also >1,000-fold attenuated for virulence in mice^49,50^. We used WT *Rp*, which does not elicit disease in WT mice^36,51^, and we also tested mutants lacking the actin-based motility factor Sca2, which is required for cell-to-cell spread and promotes dissemination in mice^52,53^. We also used a *Bt* strain lacking the motility factors BimA and MotA2^54^. C57Bl/6j mice were implanted subcutaneously with 10^6^ B16-F10 cells, which are syngeneic poorly immunogenic melanoma cells. Approximately 7 days later when tumor sizes measured ∼6 mm (width) x 6 mm (length) x 2.5 mm (depth), tumors were intratumorally injected with 10^7^ Δ*actA*Δ*inlB Lm,* Δ*bimA*Δ*motA2 Bt, sca2*::Tn *Rp*, or WT *Rp.* Each bacterial pathogen elicited a significant decrease in tumor volume as compared to vehicle PBS and promoted significantly longer survival (**Fig. 1A**). The effects were similar between the different pathogens. To determine if pathogens elicited anti-tumor effects in a different cancer indication, we measured tumor volume after intratumoral delivery of *Rp* to RMA lymphoma xenografts, which are poorly immunogenic syngeneic models of lymphoma^55^. Pathogen delivery resulted in a significant delay in tumor growth and resulted in the complete response in 5 of 19 mice (**Fig. 1B**). Among all the bacterial strains tested, no mice were euthanized due to apparent bacteremia. These data demonstrate that intratumoral delivery of phylogenetically distinct cytosolic bacterial pathogens elicits anti-cancer effects with limited/no bacterial-related toxicity.

**Fig. 1:**
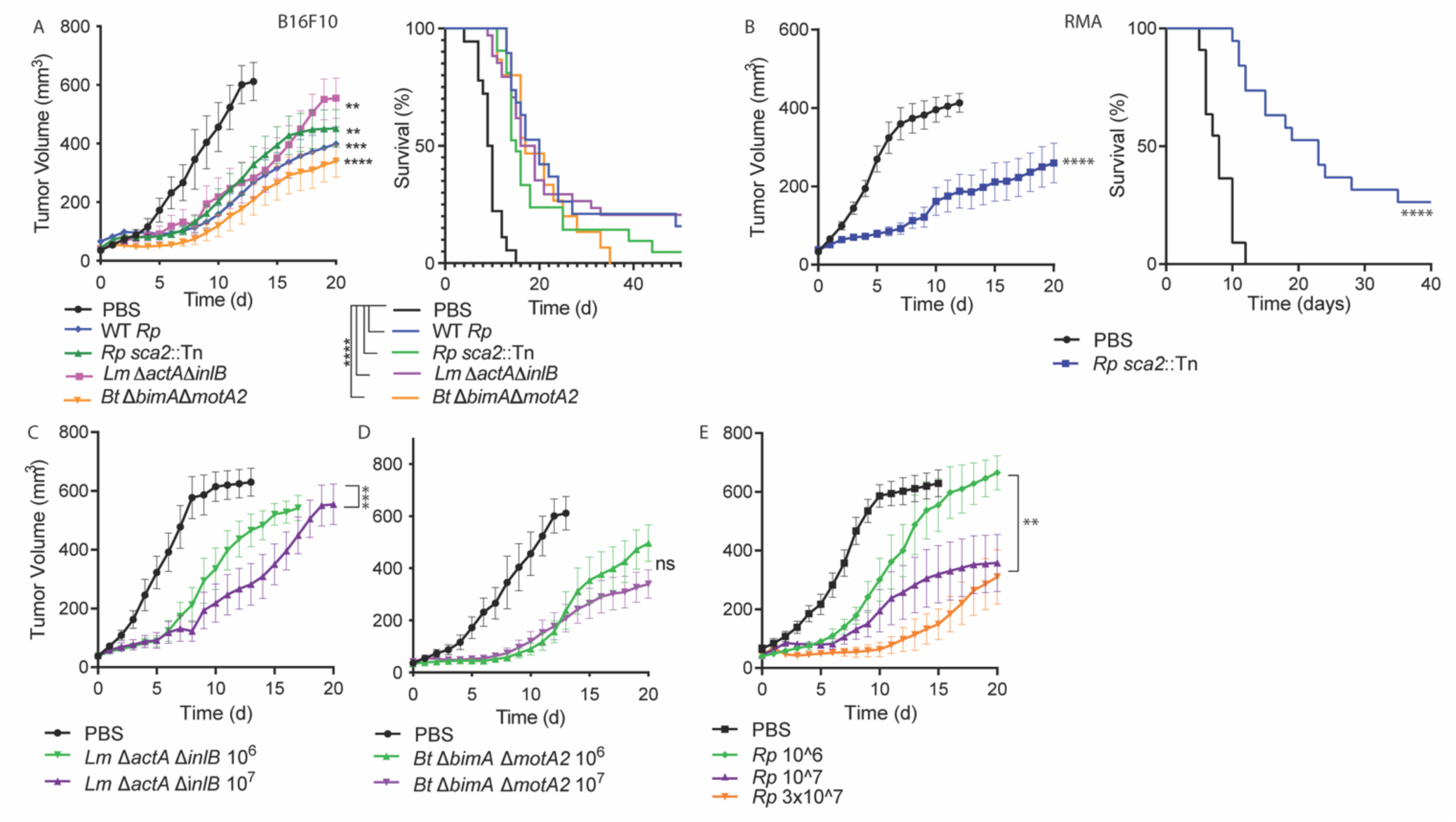
Intratumoral delivery of cytosolic bacterial pathogens elicits dose-dependent anti-tumor responses in multiple non-immunogenic murine tumor models. **a**) Tumor volume (left) and overall survival (right) of mice bearing B16-F10 tumors after therapy. Tumors measured approximately 6 x 6 x 2.5 mm in each direction and were injected with 10^7^ of the indicated bacterial species or vehicle PBS. **b**) RMA-bearing C57bl/6j mice were intratumorally injected with the indicated bacterial strain. **c-e**) Tumor volume over time of B16-F10-bearing mice treated with *Lm* (**c**), *Bt* (**d**), or *Rp* (**e**). Statistics for tumor growth used two-way ANOVA at day 20; statistics for survival used log-rank (Mantel-Cox) tests. *\*P*<0.05; \*\**P*<0.01; \*\*\**P*<0.001; \*\*\*\**P*<0.0001.

It remained unclear if the anti-tumor effects were dose dependent. We therefore examined tumor growth upon delivery of varying doses of *Lm, Bt,* and *Rp*. Delivery of 10^7^ *Lm* or *Rp* caused significantly slower tumor growth than 10^6^, while no different effects were observed with *Bt* (**Fig. 1C-E**). Delivering 3×10^7^ *Rp* did not cause significantly different responses than 10^7^ (**Fig. 1E**). These data demonstrate that the anti-tumoral effects of these pathogens are mostly dose-dependent and that 10^7^ bacteria are sufficient to maximize the anti-tumor response without eliciting toxicity. We therefore delivered 10^7^ bacteria for the remaining experiments.

### Cytosolic access of bacteria promotes the anti-tumor response

It remained unknown whether the anti-tumor effects required live bacteria to access the cytosol. We asked whether non-pathogenic *Escherichia coli* or heat-killed *Rp* elicited robust anti-tumor responses in B16-F10 tumors. Intratumoral delivery of heat-killed *Rp* or live non-pathogenic *E. coli* did not significantly delay tumor growth (**Fig. 2A**), and had a minor but significant effect on survival (**Fig. 2A**). To determine if the anti-tumor effects required access to the cytosol, we measured tumor growth after delivery of a *Lm* strain mutated for the hemolysin listeriolysin-O (LLO; encoded by the gene *hly*). *hly* mutants are unable to perforate the vacuole and are confined to membrane-bound intracellular compartments, where they do not replicate. We found that the *Lm* Δ*hly* strain did not elicit robust anti-tumor responses or improve overall survival (**Fig. 2B**). These data demonstrate that cytosolic access is necessary for eliciting a robust anti-tumor response.

**Fig 2:**
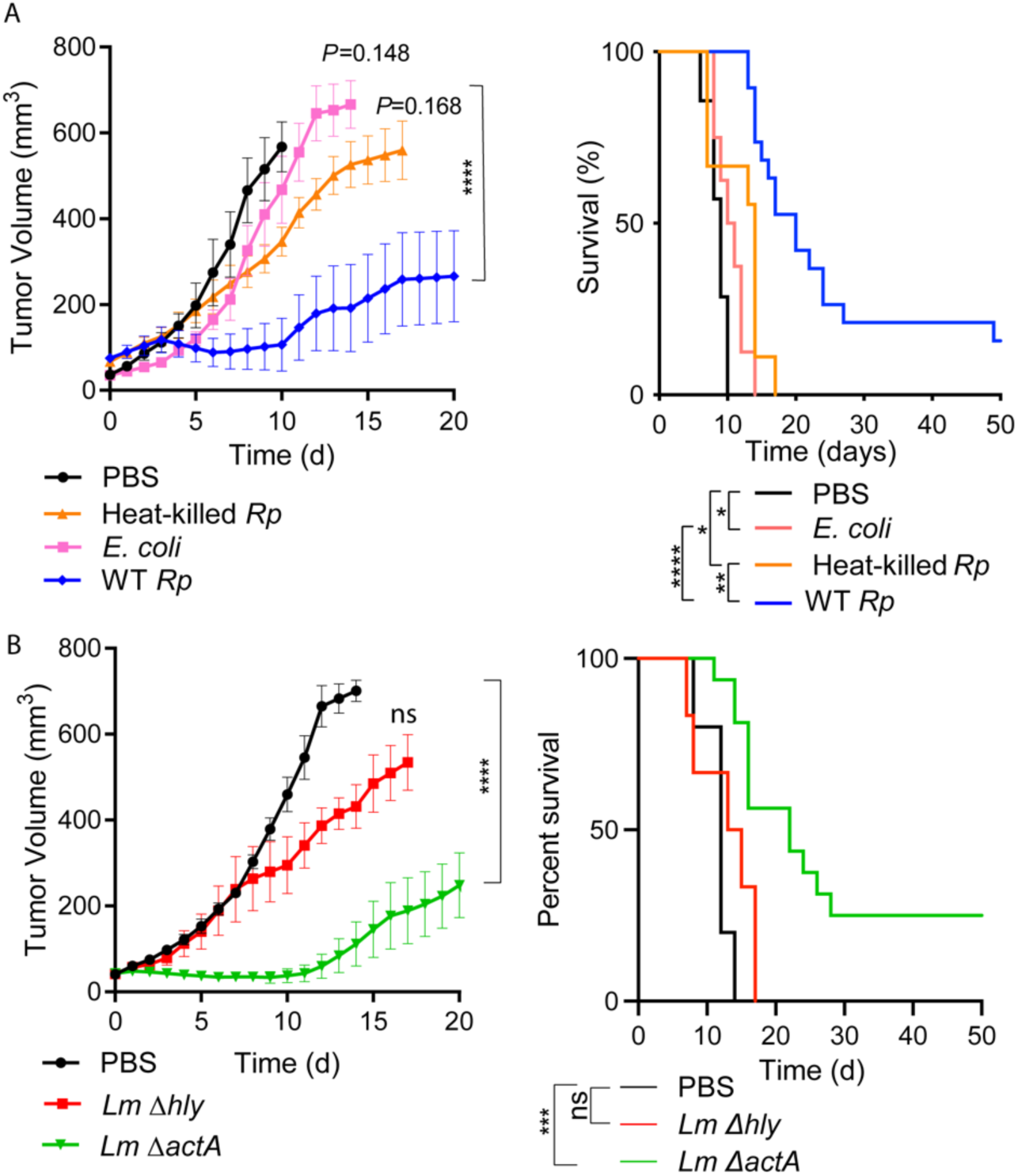
Cytosolic access is necessary for an anti-tumor response. **a-b**) Mice bearing B16-F10 tumors were subcutaneously injected with the indicated strains and tumor volume and survival were monitored over time. Statistics for tumor growth used two-way ANOVA at day 20; statistics for survival used log-rank (Mantel-Cox) tests. Statistics for tumor growth used two-way ANOVA at day 20; statistics for survival used log-rank (Mantel-Cox) tests. *\*P*<0.05; \*\**P*<0.01; \*\*\**P*<0.001; \*\*\*\**P*<0.0001. ns= not significant.

### The microbe-mediated antitumor effects are independent of cGAS/STING but require TLR signaling

It was unclear if the anti-tumor effects of *Lm/Bt/Rp* required innate immune signaling. As cytosolic access was necessary for the anti-tumor effects, we hypothesized that the anti-tumor effects were mediated via cGAS/STING. We therefore measured B16-F10 tumor volume in *Cgas*^-/-^ and *Sting^gt/gt^*mice after pathogen delivery. Contrary to our hypothesis, we observed that tumor volume after *Lm* delivery was similar between WT mice and *Cgas^-/-^*mice (**Fig. 3A**) and between WT mice and *Sting^gt/gt^* mice (**Fig. 3B**). Similar results were observed upon intratumoral delivery of *Rp* (**Fig. 3C, 3D**). No *Sting^gt/gt^*mice had complete responses **(Fig. 3E)**. It remained a possibility that cGAS signaling in the tumor cells themselves was promoting the anti-tumor response. To determine if cGAS signaling in the tumor cells contributed to the anti-tumor effects, we delivered pathogens to *Cgas*^-/-^ tumors implanted in WT and *Cgas*^-/-^ mice. The microbes elicited a similar anti-tumor effect when *Cgas^-/-^* tumor cells were implanted into either WT or *Cgas^-/-^* mice (**Fig. 3F)**, demonstrating that the antitumor response is largely independent of cGAS. Together, these data suggest that cGAS/STING only play minor roles in the anti-tumor effects mediated by cytosolic bacterial pathogens.

**Fig 3:**
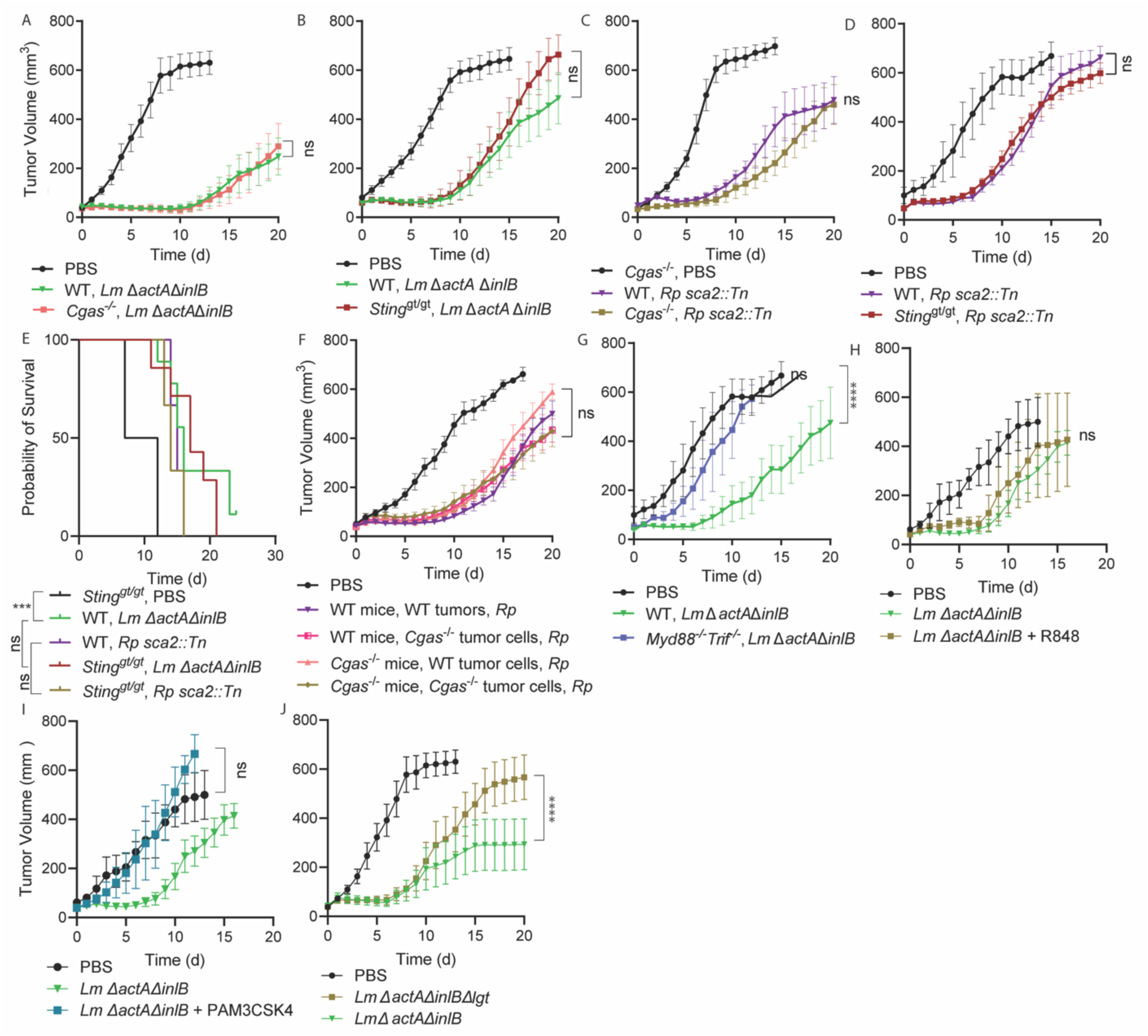
The microbe-mediated antitumor effects are independent of cGAS/STING but require TLR activation. **a-e, f-k**) The indicated strains of B16-F10-bearing mice were intratumorally administered with 10^7^ of the indicated bacterial strains and tumor volume and survival were monitored over time. **f**) The indicated strains of B16-BL6-bearing mice were intratumorally administered with 10^7^ of the indicated bacterial strains and tumor volume and survival were monitored over time. For (**j**), 10 μg of TLR7/8 agonist R848 was used and for (**k**) 10 μg of TLR2 agonist PAM3CSK4 was used. Statistics for tumor growth used two-way ANOVA at day 20; statistics for survival used log-rank (Mantel-Cox) tests. \*\*\*\**P*<0.0001. ns= not significant.

We next investigated whether other innate immune pathways were required for the microbial anti-tumor effects. Since *Lm, Rp,* and *Bt* can activate TLRs in other contexts^32,34,37,39^, and because the anti-tumor effects *M. bovis* BCG bacteria are mediated via TLR signaling^43,44,56^, we hypothesized that TLR activation contributed to the anti-tumor effects. We measured the anti-tumor responses of pathogens in *Myd88*^-/-^*Trif*^-/-^ mice and observed diminished tumor control (**Fig. 3G**), suggesting that TLR signaling is an important driver of the response. To further explore the role for TLR signaling, we hypothesized that co-administration of bacterial pathogens with small molecule TLR agonists would not dramatically enhance the anti-tumor effects. Indeed, there was no additive effect of combining the TLR7/8 agonist resiquimod (**Fig. 3H**) or the TLR2 agonist PAM3CSK4 with *Lm* (**Fig 3. I**). This provided further evidence that cytosolic pathogens elicit TLR-dependent anti-tumor responses.

We hypothesized that if the effects were mainly TLR driven, bacterial mutants deficient for lipoprotein synthesis would elicit reduced anti-tumor responses. *Lm* lipoprotein synthesis requires the phosphatidylglycerol-prolipoprotein diacylglyceryl transferase (LGT) and *lgt* mutants fail to activate TLR2 *in vivo*^57,58^. We compared the anti-tumor effects of *ΔactAΔinlB Lm* versus *ΔactAΔinlBΔlgt* and observed that strains lacking LGT had a significantly reduced anti-tumor effect (**Fig. 3J**). Taken together, these data demonstrate that, although they require cytosolic access, TLR signaling is a critical driver of the anti-tumor response to cytosolic pathogens.

### Interferons and T cells are critical for the anti-tumor effects elicited by cytosolic bacteria

We next sought to better define the role for inflammatory cytokines including interferons to the anti-tumor response elicited by cytosolic bacteria. IFN-I plays complex roles for a variety of cancer therapies and is induced by the TLR7 agonist imiquimod^59^, however IFN-I does not appear critical for the anti-tumor response elicited by BCG^56^. We observed that mice lacking the receptor for IFN-I (IFNAR) had decreased anti-tumor responses to *Lm* as compared to WT mice (**Fig. 4a**), suggesting that IFN-I contributes to the anti-tumor activities of *Lm*. We also investigated the role for IFN-γ, another pro-inflammatory cytokine that can elicit pro- or anti-tumor responses in different contexts^60^. We observed that mice lacking the receptor for IFN-γ (IFNGR) had similar anti-tumor responses as WT mice (**Fig. 4a**). As a control, we also measured tumor volume in response to S100 and in alignment with previous reports^28^, we found that it required IFNAR but not IFNGR (**Fig. 4B**). Together, these findings suggest that IFN-I but not IFN-γ contributes to the anti-tumor response elicited by *Lm*.

**Fig 4.**
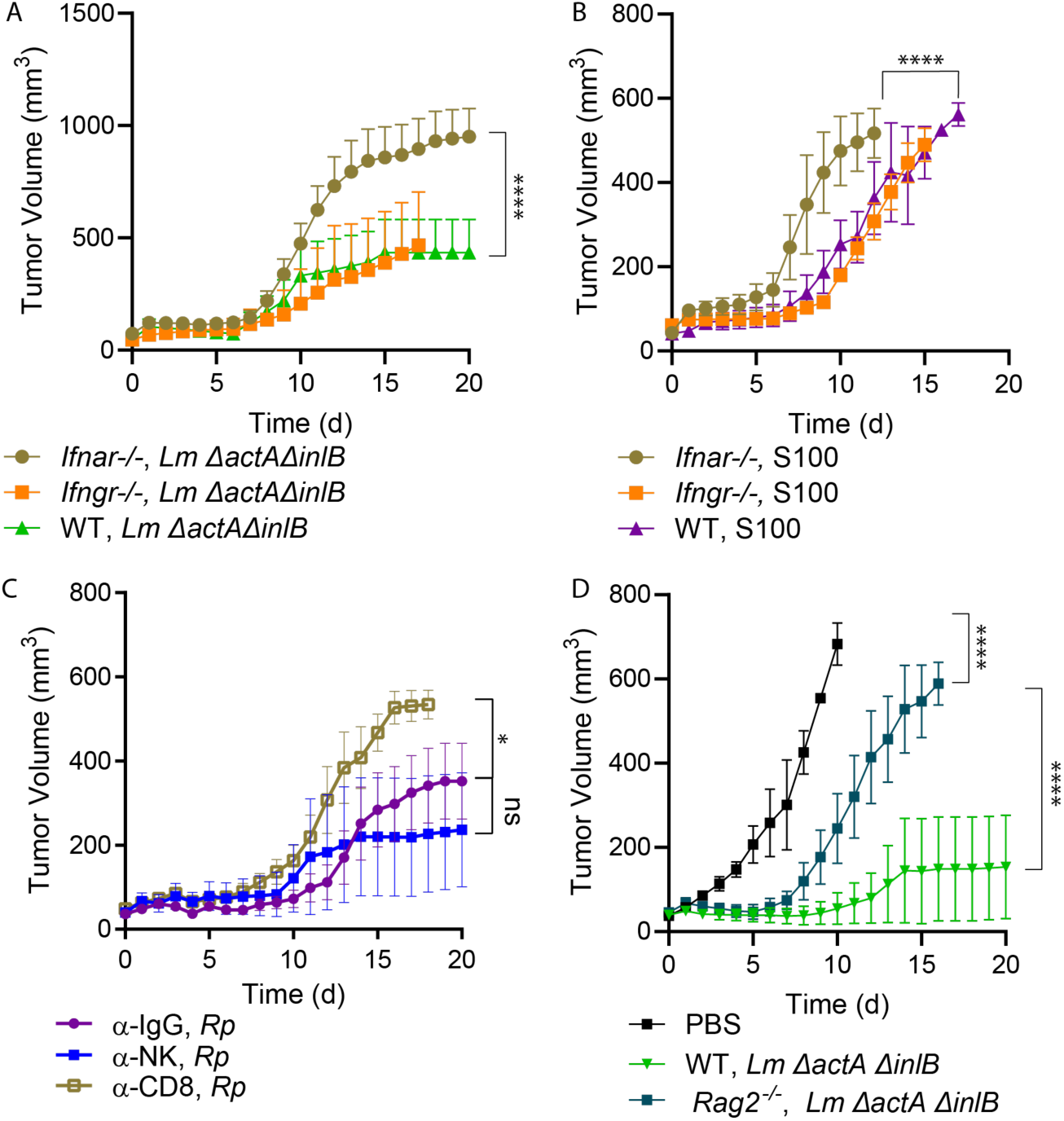
Antitumor activity of cytosolic bacteria does not require IFN-I signaling but requires CD8^+^ T cells. **a-d)** The indicated strains of B16-F10-bearing mice were intratumorally administered with 10^7^ of the indicated bacterial strains and tumor volume and survival were monitored over time. 50 μg of S100 was used and was combined with the bacteria immediately prior to injection. For (**c**) antibodies were delivered at days −2, −1, and 0. Statistics for tumor growth used two-way ANOVA at day 20; statistics for survival used log-rank (Mantel-Cox) tests. \**P*<.05, \*\*\*\**P*<0.0001. ns= not significant.

We next sought to determine the role for hematopoietic cell types including natural killer (NK) and CD8^+^ T cells in the microbe-mediated anti-tumor response. We depleted tumor-bearing WT mice of CD8^+^ T cells or NK cells and treated tumors with bacteria. Mice depleted for CD8^+^ T cells had decreased anti-tumor responses as compared to IgG control mice, while depletion of NK cells did not significantly affect the anti-tumor response (**Fig. 4C**). To further define the importance of T cells we delivered *Lm* to tumor-bearing *Rag2*^-/-^ mice, which lack all mature T and B cells. *Rag2*^-/-^ mice had a dramatically impaired ability to impede tumor growth in response to *Lm* therapy (**Fig. 4D**). Together, these experiments on cytokines and cell types demonstrate that IFN-I and T cells are critical for the anti-tumor effects elicited by cytosolic bacteria.

### STING agonists synergistically enhance the anti-tumor effects of cytosolic bacterial pathogens

Our observation that bacterial pathogens elicit TLR-dependent anti-tumor responses led us to hypothesize that their effects would be enhanced by STING agonists. We therefore evaluated the anti-tumor effects of *Lm, Bt,* and *Rp* in combination with the eukaryotic cGAS product 2’3’-cGAMP (referred here to as cGAMP) or the dithio-containing cyclic di-AMP (aka S100, ADU-S100, MIW815, ML RR-S2 CDA, or 2’3’-RR CDA)^12^. S100 binds STING with higher affinity than cGAMP and was extensively developed preclinically^12,27,28^ and underwent human trials^47,48^. To maximize the potential for observing differences between therapies, each tumor was treated with only one dose of each therapy, at d=0. Additionally, we used combinations of male and female mice that were over 18 weeks old, as we had observed that <10 week old mice respond significantly stronger to STING agonists than 18+ week old mice **(Supplemental Fig. 1).** We hypothesized that this higher threshold model would allow us to better observe differences between S100 and S100+pathogen combination therapy. Upon combining with S100, we observed striking and synergistic anti-tumor effects with *Lm* (**Fig. 5A**), *Rp,* (**Fig. 5B**) and *Bt* (**Fig. 5C**). cGAMP also dramatically enhanced the anti-tumor effects of *Lm, Rp,* or *Bt*, although to a lesser effect than S100 (**Fig. 5D-F**). Combination therapy dramatically improved overall survival with *Lm, Rp,* and *Bt* (**Fig. 5G-I**). In the case of *Lm*, combination therapy elicited complete responses in 9 of 11 mice (82%), while monotherapy with either S100 or *Lm* alone only led to complete clearance in only ∼25% of tumor-bearing mice (**Fig. 5G**). Together, these data demonstrate that the anti-tumor effects of bacteria are dramatically enhanced upon co-administration with STING agonists.

**Fig 5:**
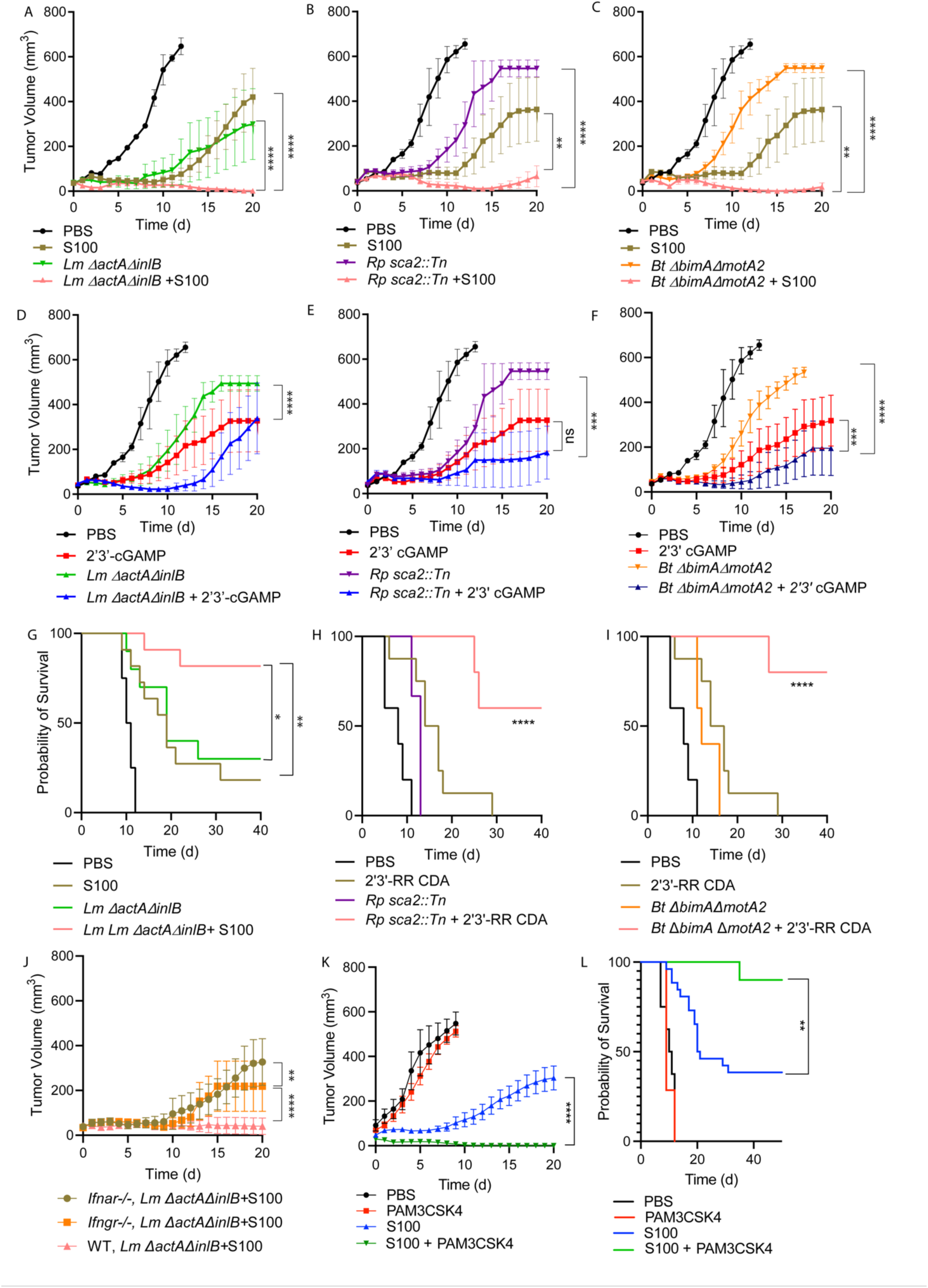
The anti-tumor effects of cytosolic bacterial pathogens synergize with cyclic dinucleotide STING agonists. B16-F10 tumor volume and survival over time after intratumoral delivery with the indicated bacterial species and CDNs. 2’3’-cGAMP and 2’3’-RR CDA was used at 50 μg / mouse. A single injection was performed for all therapies at d 0. **a-c)** *Lm, Bt,* and *Rp* alone and combined with ADU-S100 agonists and overall survival of CDA; **d-f)** *Lm, Bt,* and *Rp* alone and combined with cGAMP agonists; **g-i)** *Lm, Bt* and *Rp* alone and combined with ADU-S100 agonists overall survival**. j)** *Lm* in combination with ADU-S100 in WT, IFNAR^-/-^, and IFNGR^-/-^ mice. Statistics for tumor growth used two-way ANOVA at day 20; statistics for survival used log-rank (Mantel-Cox) tests. \*\*\*\**P*<0.0001. ns= not significant.

We previously observed that IFN-I was required for the anti-tumor effects of *Lm* and S100 therapy, however the role for interferon signaling in the combination therapy remained unknown. Indeed, we observed a significant decrease in anti-tumor efficacy from combination therapy in *Ifnar^-/-^* and *Ifngr^-/-^* mice as compared to WT mice, however, these mice still responded to combination therapy (**Fig. 5J**). This suggests that combination therapy requires IFN-I signaling but that other cytokines response are likely to also play crucial roles.

### Small molecule TLR agonists synergize with STING agonists

We next asked whether small molecular TLR agonists also synergized with STING agonists. As we observed that the production of *Lm* lipoproteins was required for the anti-tumor response, we determined whether the lipopeptide PAM3CSK4 enhanced the anti-tumor effects of S100. Similar to the bacterial pathogens, we observed that S100 anti-tumor activity was dramatically enhanced by the addition of PAM3CSK4 (**Fig. 5K**), as was survival (**Fig. 5L**). Notably, unlike *Lm*/*Rp*/*Bt*, PAM3CSK4 had no anti-tumor effects on its own, in alignment with previous observations^61^, suggesting that the bacterial pathogens activate stronger anti-tumor responses than TLR2 agonists alone.

### Mice that clear initial tumors after microbial therapy have increased immunity to tumor cell rechallenge

It remained unknown whether mice that received therapy and cleared the initial tumor had a long-lived adaptive immune response against cancer. We therefore next examined if mice that rejected tumors after microbial treatment developed tumors after re-administration of the same tumor cells >40 days later. 6 of 8 mice that cleared initial B16-F10 tumors by *Lm* and 4 of 8 mice that cleared tumors by S100 rejected tumor rechallenges, whereas tumors expanded in naïve mice (**Fig. 6A**). Among mice that cleared initial tumors after combination pathogen + CDN therapy, 6 of 9 mice that cleared initial tumors after combinational therapy were also resistant to tumor cell rechallenge (**Fig. 6a**). This suggested that bacterial therapy alone or in combination with STING agonists elicits long-lasting protection against cancer.

**Fig. 6:**
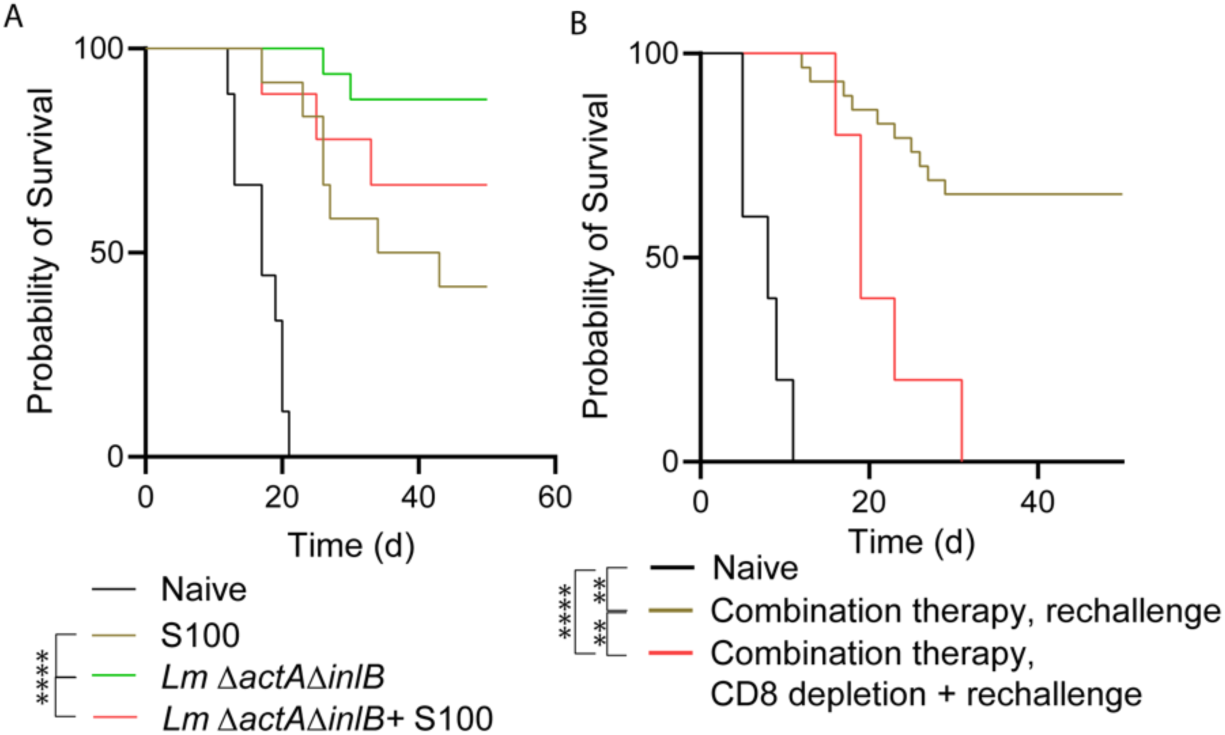
Mice that clear initial tumors after microbial therapy have increased immunity to tumor cell rechallenge. **a-b**) Survival of mice after rechallenge with B16-F10 tumors. 10^6^ B16-F10 cells were implanted subcutaneously into mice that cleared initial tumors after intratumoral delivery of the indicated therapies. Survival of mice after rechallenge with B16 following CD8 T cell depletion. 10^6^ B16 cells were implanted subcutaneously into mice that cleared initial tumors after intratumoral delivery with various.

To determine if this protective immunity was T cell dependent, we depleted CD8^+^ T cells in mice that cleared the initial tumor and then rechallenged them with 10^6^ B16-F10 tumor cells in the opposite flank. Mice depleted for CD8 T cells demonstrated a decreased ability to reject the tumor cell rechallenge (**Fig. 6b**). These findings demonstrate that intratumoral delivery of cytosolic bacterial pathogens and combinational therapy of pathogens with STING agonists elicits long-lasting protective immune responses against cancer that require CD8^+^ T cells.

## Discussion

Bacteria have been used to treat cancer for over 100 years and they are the first examples of immunotherapy^6^. Yet the anti-tumor potential for cytosolic bacterial pathogens, which interface with a distinct set of PRRs in the cytosol, has remained unknown. Here, we find that phylogenetically distinct species of Gram-positive and Gram-negative cytosol-dwelling bacterial pathogens elicit anti-tumor responses in mice. The anti-tumor responses require access to the cytosol, but are largely independent of cGAS/STING, and instead require TLR signaling. Strikingly, we find that combining cytosolic pathogens with STING agonists elicits a synergistic anti-tumor effect that clears injected tumors with a high frequency and elicits a long-lasting CD8^+^ T cell response against cancer. This strategy is highly effective even with established, poorly immunogenic B16-F10 melanomas in male and female mice aged >18 weeks old. We propose that the co-activation of STING and TLRs is a robust strategy for designing the next generations of microbial and small molecule-based innate immune agonist therapies.

Our results suggest that live cytosolic bacteria pathogens elicit superior anti-tumor responses as compared to heat-killed bacteria, non-pathogenic bacteria, and small molecule TLR2 agonists. This could be due to bacterial co-activation of multiple PRRs or because bacterial growth in the cytosol increases the number of innate immune pathways activated over time. Bacterial vectors are being tested clinically as vehicles to deliver STING agonists, including a non-pathogenic *E. Coli* Nissle strain engineered to express cyclic di-AMP in the tumor. Intratumoral injection of this strain to B16-F10 tumor-bearing mice induces IFN-I production and reduces tumor growth^62^. This study also found that *E. coli* activate TLRs *in vitro*. In a phase I clinical trial (NCT04167137), this cyclic di-AMP-expressing *E. coli* strain was delivered intratumorally as monotherapy or in combination with Atezolizumab and demonstrated safety and cytokine production^63,64^. Based on our findings, we speculate that this approach may be activating TLR and STING pathways, although the magnitude of these effects when compared to a cocktail of STING agonists and *E. coli* is unknown. As we found that S100 elicits superior responses to cGAMP, which has higher affinity for STING than CDA, these strains would likely be improved if they were able to secrete agonists with enhanced binding affinity for STING. Another novel bacterial-based immunotherapy is an attenuated *Salmonella* Typhimurium strain (STACT) that carries an inhibitor of TREX-1, the exonuclease that prevents activation of STING by degrading cytosolic DNA. Pre-clinical work found that intravenously delivery caused tumor colonization, tumor regression, and immunity to rechallenge^45,65^. Such microbial-based cancer therapies are advantageous as they can be administrated systemically and thus can target tumors throughout the body. However, in these studies the role for co-activation of STING and TLRs has not been explicitly appreciated, and based on our work we hypothesize that a lynchpin for their efficacy is robust activation of these pathways.

Cytokines including interferons play multifaceted roles in cancer, in which acute therapeutic activation of STING requires IFN-I signaling for a proper anti-tumor response^28,59,66^. IFN-I promotes the ability of dendritic cells to cross-present antigen to T cells, and activation^66,67^, and CD8a dendritic cells are required to spontaneously prime tumor-specific CD8+ T cells^68,69^. IFN-I is induced by the TLR7 agonist imiquimod^70^, however IFN-I does not appear critical for the anti-tumor response elicited by BCG^56^. In this study we observed that IFN-I and IFN-γ are required for the anti-tumor response to bacterial pathogens alone but are mostly dispensable for STING+TLR agonist combination therapy. Other studies that observed STING and TLR agonist synergy for cancer therapy reported that IFN-I and other cytokines including IL-12 are synergistically produced *in vitro* and *in vivo;* however, cytokine neutralization studies *in vivo* or studies using mutant mice lacking these cytokine signaling pathways are lacking^71–78^. Thus, our work suggests for the first time that despite the fact that STING+TLR combination therapy synergistically produces pro-inflammatory cytokines, this response may not actually be essential for the anti-tumor response to this therapy.

Spontaneous T cell development against tumors have been shown to improve patients’ overall prognosis, and STING agonists elicit long-lasting T cell responses in preclinical models^12,28,79^. In alignment with this, we find that *Rag2*^-/-^ mice have decreased antitumor responses, and CD8 T cell depletion reduces immunity against tumor rechallenge. One challenge with STING+TLR combination therapy is balancing an anti-tumor response with excessive inflammation can result in inhibitory, apoptotic effects for infiltrating T cells^80^. Previous work with STING agonists elegantly demonstrated that 50 ug of intratumoral delivery of S100 elicits a strong initial anti-tumor response but also a long-lasting memory response, while higher doses can cause T cell apoptosis to the detriment of the immune response^80^. Our data shows that the combinatorial effects of pathogens with STING agonists are highly potent in reducing tumor size, and the cured mice have long-lasting protection that is similar to S100 therapy alone. Future studies that more closely investigate the T cell response to STING+TLR agonist therapy are warranted to identify the optimal drug dosage combinations for eliciting both an aggressive anti-tumor response paired with a strong memory response.

This study focused on intratumoral deliveries as a robust methodology that allowed us to finely discriminate between the efficacy of certain therapies. However, this methodology does not robustly model metastatic cancer, in which systemic therapy is more likely to cause tumor regression across many distal tumors. Thus, developing STING+TLR combination therapies for systemic delivery is a critical future direction of this work. One challenge towards developing systemic therapies with innate immune agonists, however, is that STING and TLRs are widely expressed on many resident tissue cell types, including endothelial cells, macrophages, and monocytes^81^. This hurdle will need to be overcome by specifically targeting tumors, perhaps through tumor-targeting bacteria or small molecules that are activated preferentially in tumors.

## Methods

### Animal maintenance

Animal research using mice was conducted under a protocol approved by the UC Berkeley Institutional Animal Care and Use Committee (IACUC) in compliance with the Animal Welfare Act and other federal statutes relating to animals and experiments using animals (Welch lab animal use protocol AUP-2016-02-8426). The UC Berkeley IACUC is fully accredited by the Association for the Assessment and Accreditation of Laboratory Animal Care International and adheres to the principles of the Guide for the Care and use of Laboratory Animals. Infections were performed in a biosafety level 2 facility and all animals were maintained at the UC Berkeley campus. All mice were healthy at the time of tumor delivery and were housed in microisolator cages and provided chow and water. No mice were administered antibiotics or maintained on water with antibiotics.

Mice were between 8 and 24 weeks old at the time of tumor delivery and all mice were of the C57BL/6J background. Mice were selected for experiments based on their availability and both male and female mice were used in experiments. Initial sample sizes were based on availability of mice, which were approximately 5 mice per group and a minimum of 3 mice per group. Therapeutic treatments were assigned in an effort to divide each therapy into as many cages as possible and with an even number of male/female mice. Mice were euthanized if tumor diameter exceeded 15 mm in any direction. After the first experiment, a Power Analysis was conducted to determine subsequent group sizes.

### Tumor xenografts and intratumoral deliveries

B16-F10, B16-Bl6, and RMA cells were grown *in vitro* in DMEM (Gibco 11965-092) supplemented with 10% fetal bovine serum (FBS, Corning 35-010-CV). Prior to injection, cells were trypsinized, counted, washed twice with sterile PBS, and resuspended at 1.5×10^6^ cells/100 ml. Mice were shaved on their right hind flank and injected subcutaneously with tumor cells in 100 ml volumes. Tumor size was monitored by measuring the length, width, and height of each tumor using calipers, where *V* = (length × width × height)*3.1415/6, as described previously^55^. Tumors were injected when they had reached the approximate dimensions of 6 x 6 x 2.5 mm. On days when tumors were not measured, the growth in tumor volume was calculated by taking the difference between tumor volumes at adjacent time points.

### Preparation of bacteria

*Rp* strain Portsmouth was originally obtained from Christopher Paddock (Centers for Disease Control and Prevention). Bacteria were amplified by infecting confluent T175 flasks of female African green monkey kidney epithelial Vero cells authenticated by mass spectrometry. WT and *sca2* mutant *Rp* stocks were purified and quantified as described^82–85^. For mouse infections, *Rp* was prepared by diluting 30%-prep bacteria into sterile PBS on ice, centrifuging the bacteria at 12,000 x G for 1 min (Eppendorf 5430 centrifuge), and resuspended in cold sterile PBS to the desired concentration (either 10^7^ PFU/ 50 μl or 10^6^ PFU/50 μl). Bacterial suspensions were kept on ice during injections. Mice were scruffed and 50 µl of bacterial suspensions were injected using 30.5-gauge needles into palpable tumors. Body temperatures were monitored using a rodent rectal thermometer (BrainTree Scientific, RET-3). CD8^+^ T cells were depleted by injecting mice IP with 160 µg of α-CD8b.2 (Leinco C2832) on days −2 and −1 prior to infection (320 µg total per mouse). NK cells were depleted by injecting mice IP with 200 µg PK136 antibody on days −2 and −1 prior to infection. For control experiments, 100 µg of control IgG antibody (Jackson, 012-000-003) was delivered IP at days −2 and −1. After infection, all mice in this study were monitored daily for clinical signs of disease, such as hunched posture, lethargy, or scruffed fur.

*Lm* and *Bt* were prepared by inoculating 2 ml liquid brain heart infusion (BHI) media into 14 ml conical tubes and growing the bacteria for 20 h shaking at a slant at 37° C. Bacteria were then diluted 1:40 into 2 ml fresh BHI and grown for 2 h. OD_600_ for each sample was measured, bacteria were centrifuged and washed once with sterile room temperature PBS, and resuspended in PBS to a concentration of 10^7^ or 10^6^/ 50 μl. *Lm* and *Bt* were kept at room temperature prior to injection and delivered intratumorally using 30.5 gauge needles. Bacteria were serially diluted and plated on LB plates to verify the inoculum.

### Mouse genotyping

*Sting^gt/gt^* and *Cgas^-/-^* mice were generated at UC Berkeley, as previously described^23,86^. *Ifnar^-/-^* ^87^*, Ifngr^-/-^*^88^, and *Rag2^-/-^* mice were previously described and obtained from Jackson Labs. C57BL/6J WT mice were originally obtained from Jackson Laboratories. For genotyping, ear clips were boiled for 15 min in 60 µl of 25 mM NaOH, quenched with 10 µl tris-HCl pH 5.5, and 2 µl of lysate was used for PCR using SapphireAMP (Takara, RR350) and primers specific for each gene. Mice were genotyped using these primers: *Cgas* F: ACTGGGAATCCAGCTTTTCACT; *Cgas* R: TGGGGTCAGAGGAAATCAGC; *Sting* F: GATCCGAATGTTCAATCAGC; *Sting* R: CGATTCTTGATGCCAGCAC; *Ifnar* forward (F): CAACATACTACAACGACCAAGTGTG; *Ifnar* WT reverse (R): AACAAACCCCCAAACCCCAG; *Ifnar* mutant R: ATCTGGACGAAGAGCATCAGG;

### Deriving bone marrow macrophages

To obtain bone marrow, male or female mice were euthanized, and femurs, tibias, and fibulas were excised. Connective tissue was removed, and the bones were sterilized with 70% ethanol. Bones were washed with BMDM media (20% FBS, 1% sodium pyruvate, 0.1% β-mercaptoethanol, 10% conditioned supernatant from 3T3 fibroblasts, in Gibco DMEM containing glucose and 100 U/ml penicillin and 100 ug/ml streptomycin) and ground using a mortar and pestle. Bone homogenate was passed through a 70 μm nylon Corning Falcon cell strainer (Thermo Fisher Scientific, 08-771-2) to remove particulates. Filtrates were centrifuged in an Eppendorf 5810R at 1,200 RPM (290 x G) for 8 min, supernatant was aspirated, and the remaining pellet was resuspended in BMDM media. Cells were then plated in non-TC-treated 15 cm petri dishes (at a ratio of 10 dishes per 2 femurs/tibias) in 30 ml BMDM media and incubated at 37° C. An additional 30 ml was added 3 d later. At 7 d the media was aspirated, and cells were incubated at 4°C with 15 ml cold PBS (Gibco, 10010-023) for 10 min. BMDMs were then scraped from the plate, collected in a 50 ml conical tube, and centrifuged at 1,200 RPM (290 x G) for 5 min. The PBS was then aspirated, and cells were resuspended in BMDM media with 30% FBS and 10% DMSO at 10^7^ cells/ml. 1 ml aliquots were stored in liquid nitrogen.

### Infections *in vitro*

To plate cells for infection, aliquots of BMDMs were thawed on ice, diluted into 9 ml of DMEM, centrifuged in an Eppendorf 5810R at 1,200 RPM (290 x G) for 5 minutes, and the pellet was resuspended in 10 ml BMDM media without antibiotics. The number of cells was counted using Trypan blue (Sigma, T8154) and a hemocytometer (Bright-Line), and 5 x 10^5^ cells were plated into 24-well plates. Approximately 16 h later, 30% prep *Rp* were thawed on ice and diluted into fresh BMDM media to the desired concentration (either 10^6^ PFU/ml or 2×10^5^ PFU/ml). Media was then aspirated from the BMDMs, replaced with 500 µl media containing *Rp,* and plates were spun at 300 G for 5 min in an Eppendorf 5810R. Infected cells were then incubated in a humidified CEDCO 1600 incubator set to 33°C and 5% CO_2_ for the duration of the experiment. For treatments with recombinant mouse IFN-β, IFN-β (PBL, 12405-1) was added directly to infected cells immediately after spinfection.

For infections with *Lm,* cultures of *Lm* strain 10403S (originally obtained from Dr. Dan Portnoy, UC Berkeley) were grown in 2 ml sterile-filtered BHI shaking at 37° to stationary phase (∼16 h). Cultures were centrifuged at 20,000 x G (Eppendorf 5430), the pellet was resuspended in sterile PBS and diluted 100-fold in PBS. 10 µl of the diluted bacteria were then added to each well of a 24-well plate of BMDMs that were plated ∼16 h prior to infections at 5×10^5^ cells/well. Bacteria were also plated out onto Luria Broth agarose plates to determine the titer, which was determined to be ∼5 x 10^5^ bacteria / 10 µl, for an MOI of 1 (based on the ratio of bacteria in culture to number of BMDMs). Infected cells were incubated in a humidified 37° incubator with 5% CO_2_. 25 µg of gentamicin (Gibco 15710-064) was added to each well (final concentration 50 µg/ml) at 1 hpi. At 30 mpi, 2, 5, and 8 hpi, the supernatant was aspirated from infected cells, and cells were washed twice with sterile milli-Q water. Infected BMDMs were then lysed with 1 ml sterile water by repeated pipetting and scraping of the well. Lysates were then serially diluted and plated on LB agar plates, incubated at 37° overnight, and CFU were counted at ∼20 h later.

### In vitro assays

For LDH assays, 60 µl of supernatant from wells containing BMDMs was collected into 96-well plates. 60 µl of LDH buffer was then added. LDH buffer contained: 3 µl of “INT” solution containing 2 mg/ml tetrazolium salt (Sigma I8377) in PBS; 3 µl of “DIA” solution containing 13.5 units/ml diaphorase (Sigma, D5540), 3 mg/ml β-nicotinaminde adenine dinucleotide hydrate (Sigma, N3014), 0.03% BSA, and 1.2% sucrose; 34 µl PBS with 0.5% BSA; and 20 µl solution containing 36 mg/ml lithium lactate in 10 mM Tris HCl pH 8.5 (Sigma L2250). Supernatant from uninfected cells and from cells completely lysed with 1% triton X-100 (final concentration) were used as controls. Reactions were incubated at room temperature for 20 min prior to reading at 490 nm using an Infinite F200 Pro plate reader (Tecan). Values for uninfected cells were subtracted from the experimental values, divided by the difference of triton-lysed and uninfected cells, and multiplied by 100 to obtain percent lysis. Each experiment was performed and averaged between technical duplicates and biological triplicates.

For the IFN-I bioassay, 5 x 10^4^ 3T3 cells containing an interferon-sensitive response element (ISRE) fused to luciferase were plated per well into 96-well white-bottom plates (Greneir 655083) in DMEM containing 10% FBS, 100 U/ml penicillin and 100 µg/ml streptomycin. Media was replaced 24 h later and confluent cells were treated with 2 µl of supernatant harvested from BMDM experiments. Media was removed 4 h later and cells were lysed with 40 µl TNT lysis buffer (20 mM Tris, pH 8, 200 mM NaCl, 1% triton-100). Lysates were then injected with 40 µl firefly luciferin substrate (Biosynth) and luminescence was measured using a SpectraMax L plate reader (Molecular Devices).

### Statistical analysis

Statistical parameters and significance are reported in the figure legends. For tumor growth, comparisons were made using two-way ANOVAs. For survival, log-rank (Mantel-Cox) tests were used. For comparing two sets of data, including for bacterial growth curves, a two-tailed Student’s T test was performed for each time point. For comparing multiple data sets, including host cell death and IFN-I assays, a one-way ANOVA with multiple comparisons with Tukey post-hoc test was used for normal distributions. Data are determined to be statistically significant when *P*<0.05. For tumor growth curves, data are the means and error bars represent the standard error of the mean (SEM). In bar graphs, all data points are shown which represent biological replicates, and error bars represent standard deviation (SD). Asterisks denote statistical significance as: *, *P*<0.05; **, *P*<0.01; ***, *P*<0.001; ****, *P*<0.0001. All other graphical representations are described in the Figure legends. Statistical analyses were performed using GraphPad PRISM V9.

## Data availability

*Rp* strains were authenticated by whole genome sequencing and are available in the NCBI Trace and Short-Read Archive; Sequence Read Archive (SRA), accession number SRX4401164.

## Additional Information

Correspondence and requests for materials should be addressed to T.P.B.

## Acknowledgements

C.J.N. was supported by NIH fellowship F31CA228381. T.P.B. was supported in part by ACS Seed Grant 129801-IRG-16-187-13-IRG from the American Cancer Society

## Author contributions

M.D. and T.P.B. designed, performed, and analyzed experiments. N.W., T.T.V., and C.J.N. contributed to performing experiments. T.P.B. and M.D. wrote the original draft of this manuscript. Critical reading and edits were provided M.D., N.W., C.J.N., and T.T.V. Supervision was provided by T.P.B.

