## Supplemental Figures for "Cytosolic bacterial pathogens activate TLR pathways in tumors that synergistically enhance STING agonist cancer therapies"

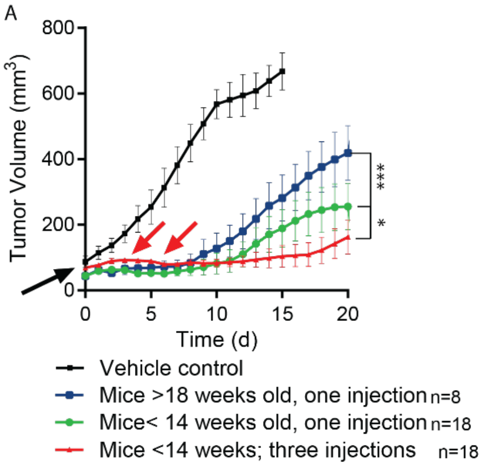


**Supplemental Fig 1. Younger mice have a more robust response to immunotherapy than older mice.** B16-F10-bearing mice were intratumorally administered PBS or S100 at >18 (*n=8)* and <14 *(n=18)* weeks of age. A second group of <14 week old mice *(n=18)* received 3 injections at days 0, 3, and 6. Statistics for tumor growth used two-way ANOVA; statistics for survival used log-rank (Mantel-Cox) tests. *P<0.05, ***P<0.001.

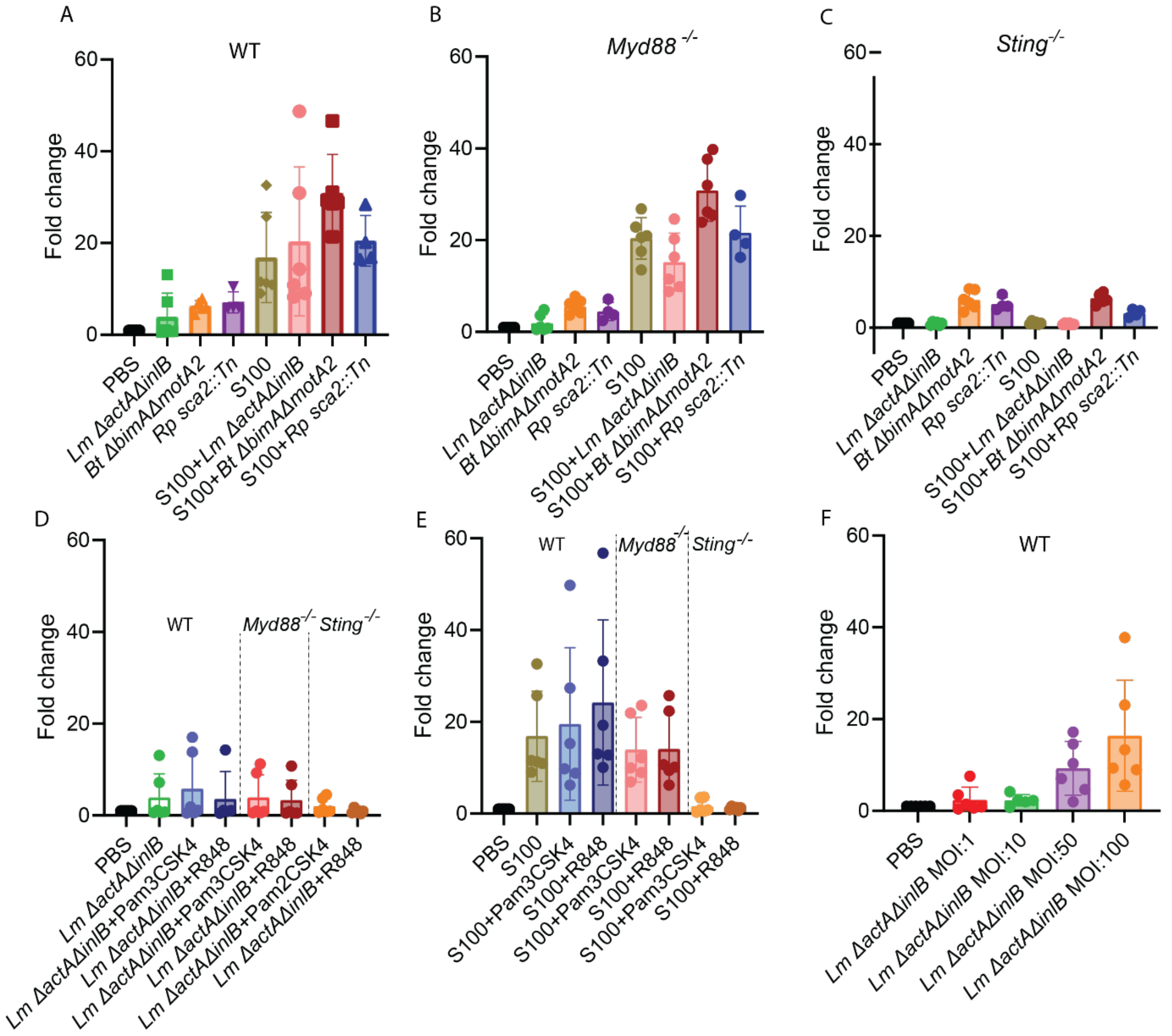

**Supplemental Fig. 2. IFN-I production is higher in BMDMs treated with pathogen with S100.** A-f) IFN-I abundance produced by murine bone marrow derived macrophages (BMDMs) treated with pathogens or small molecules after 7 hours, *n=3*. WT **a)**, *Myd88-/-* **b)**, and *Sting-/-* **c)** were infected with *Lm, Bt,* and *Rp* with or without S100. **d)** WT, *Myd88-/-*, or *Sting-/-* BMDMs treated with *Lm* with or without PAM3CSK4 or R848. **e)** WT, *Myd88-/-*, or *Sting-/-* BMDMs treated with S100 with or without PAM3CSK4 or R848. **f)** WT BMDMs treated with *Lm* at 1, 10, 50, and 100 MOI.
